# RNA virus discovery sheds light on the virome of a major vineyard pest, the European grapevine moth (*Lobesia botrana*)

**DOI:** 10.1101/2024.12.13.628328

**Authors:** Humberto Debat, Sebastian Gomez-Talquenca, Nicolas Bejerman

**Affiliations:** Instituto de Patología Vegetal – Centro de Investigaciones Agropecuarias – Instituto Nacional de Tecnología Agropecuaria (IPAVE-CIAP-INTA), Camino 60 Cuadras Km 5,5 (X5020ICA), Córdoba, Argentina; Unidad de Fitopatología y Modelización Agrícola – Consejo Nacional de Investigaciones Científicas y Técnicas (UFYMA-CONICET)., Camino 60 Cuadras Km 5,5 (X5020ICA), Córdoba, Argentina; Estación Experimental Agropecuaria Mendoza – Instituto Nacional de Tecnología Agropecuaria (EEA-Mendoza-INTA), San Martín 3853, Luján de Cuyo, Mendoza, 5534, Argentina

## Abstract

The European grapevine moth (*Lobesia botrana*) poses a significant threat to vineyards worldwide, causing extensive economic losses. While its ecological interactions and control strategies are well-studied, its associated viral diversity remains unexplored. Here, we employ high-throughput sequencing data mining to comprehensively characterize the *L. botrana* virome, revealing novel and diverse RNA viruses. We characterized four new viral members belonging to distinct families, with evolutionary cues of cypoviruses (*Reoviridae*), sobemo-like viruses (*Solemoviridae*), phasmaviruses (*Phasmaviridae*), and carmotetraviruses (*Carmotetraviridae*). Phylogenetic analysis of the cypoviruses places them within the genus in affinity with other moth viruses, suggesting a potential ecological roles and host range within Lepidoptera. The bi-segmented and highly divergent sobemo-like virus showed a distinctive evolutionary trajectory of its encoding proteins at the periphery of recently reported invertebrate Sobelivirales. Notably, the presence of a novel phasmavirus, typically associated with mosquitoes, expands the known host range and diversity of this family to moths. Furthermore, the identification of a carmotetravirus branching in the same cluster of providence virus, a lepidopteran virus which replicates in plants, raises questions regarding the biological significance of this moth virus and grapevine. We further explored viral sequences in several publicly available transcriptomic datasets of the moth, indicating potential prevalence across distinct conditions. These results underscore the existence of a complex virome within *L. botrana* and lays the foundation for future studies investigating the ecological roles, evolutionary dynamics, and potential biocontrol applications of these viruses on the *L. botrana*—vineyard ecosystem.

## Introduction

The European grapevine moth, *Lobesia botrana*, poses a pervasive threat to vineyards globally, exerting substantial economic effects for grape growers [1]. Despite extensive research on its ecology, behavior, and management strategies [2–4], a significant gap persists in our knowledge on the *L. botrana* virome. This knowledge gap hinders a comprehensive understanding of *L. botrana* population dynamics and its susceptibility to environmental stressors. Viral infection has demonstrable effects on arthropod biology, influencing fitness, behavior, and resistance to environmental changes [5, 6]. In agricultural ecosystems, these interactions can profoundly impact productivity, pest dynamics and the efficacy of control measures [7, 8]. Consequently, uncovering the composition and diversity of the *L. botrana* virome is essential to understand this plague and design more effective pest management strategies.

In this line, we initiated here an extensive exploration of the *L. botrana* virome through the integration of high-throughput sequencing (HTS) outputs. This *in silico* strategy allowed us to surpass the limitations of traditional virus isolation techniques, capturing an unbiassed viral repertoire of invertebrates, including both known and potentially novel viral lineages [9, 10]. Our investigation focused on characterizing the *L. botrana* virome to comprehensively define the taxonomic composition and diversity of novel viral lineages, unveiling the hidden viral counterpart shaping the moth’s biology. By shedding light on the previously unseen viral dimension of *L. botrana* biology, we aimed to provide valuable insights to foster future studies on the ecological roles of viruses within this significant agricultural pest. Additionally, a glimpse of the *L. botrana* virome contributes to a more comprehensive understanding of insect-virus interactions within agricultural ecosystems, potentially impacting broader ecological and evolutionary studies, valuable for the development of novel, virus-based control strategies. This study introduces a pivotal step towards comprehending a crucial facet of this economically important pest and contributing to a more sustainable future for vineyards.

## Materials and Methods

### Database selection and High-Throughput Sequencing (HTS) Library processing

To characterize the *L. botrana* virome, publicly available metatranscriptomic RNA-Seq datasets from *L. botrana*individuals collected from diverse geographic regions were downloaded from the NCBI Sequence Read Archive (SRA). *L. botrana* RNA was prepared and sequenced as described in [11], [12] and Reineke et al. (PRJNA910346, unpublished). The *Lobesia botrana* individuals sampled and used for virus discovery correspond to both larvae and adults, with RNA extracted from whole individuals or from pheromone glands, including larvae, hosted on *Vitis vinifera* cv Cabernet Sauvignon or cv Riesling during flowering or veraison developmental stage exposed to elevated (ca. 480 ppm) or ambient (ca. 400 ppm) CO2 levels, or individuals susceptible or resistant to insecticides, and from Turkey or Germany. The datasets included in this work are available in the NCBI SRA archive at https://www.ncbi.nlm.nih.gov/sra/?term=txid209534[Organism:noexp] and described in **Table 1**.

### Bioinformatics analysis, sequence assembly and virus identification

Virus discovery procedures were conducted following established methodologies [13, 14]. Briefly, raw nucleotide sequence reads from each *L. botrana* SRA experiment were retrieved from their respective NCBI BioProjects (**Table 1**). The datasets underwent preprocessing, including trimming and filtering, using the Trimmomatic v0.40 tool, accessed via http://www.usadellab.org/cms/?page=trimmomatic on October 30, 2024. Standard parameters were employed, apart from raising the quality required from 20 to 30 (initial ILLUMINACLIP step, sliding window trimming, average quality required = 30). The resulting reads were assembled de novo using rnaSPAdes with standard parameters on the Galaxy server (https://usegalaxy.org/), accessed on October 30, 2024. Subsequently, transcripts obtained from the *de novo* transcriptome assembly underwent bulk local BLASTX searches (E-value < 1e−5) against the complete NR release of viral protein sequences which was retrieved from https://www.ncbi.nlm.nih.gov/protein/?term=txid10239[Organism:exp], accessed on October 30, 2024. Resulting viral sequence hits from each dataset were thoroughly examined. Tentative virus-like contigs were curated, extended, and/or confirmed through iterative mapping of filtered reads from each SRA library. This iterative strategy involved extracting a subset of reads related to the query contig, utilizing the retrieved reads to extend the contig, and repeating the process iteratively using the extended sequence as the new query. The extended and polished transcripts were subsequently reassembled using the Geneious v8.1.9 alignment tool (Biomatters Ltd., Boston, MA, USA) with high sensitivity parameters.

### Bioinformatics characterization of novel viral genomes

Open reading frames (ORFs) were predicted using ORFfinder with a minimal ORF length of 150 nt and genetic code 1 (https://www.ncbi.nlm.nih.gov/orffinder/, accessed on October 30, 2024). The functional domains and architecture of translated gene products were determined using InterPro (https://www.ebi.ac.uk/interpro/search/sequence-search, accessed on October 30, 2024) and the NCBI Conserved domain database-CDD v3.20 (https://www.ncbi.nlm.nih.gov/Structure/cdd/wrpsb.cgi, accessed on October 30, 2024) with an e-value threshold of 0.01. Additionally, HHPred and HHBlits, as implemented in https://toolkit.tuebingen.mpg.de/#/tools/ (accessed on October 30, 2024), were employed to complement the annotation of divergent predicted proteins using hidden Markov models. Transmembrane domains were predicted using the TMHMM version 2.0 tool (http://www.cbs.dtu.dk/services/TMHMM/, accessed on October 30, 2024). The predicted proteins were then subjected to NCBI-BLASTP web searches against the non-redundant protein sequences (nr) database to filter out any endogenous virus-like sequences that did not show virus protein as the best hit. Phylogenetic analysis based on the predicted polymerase protein of all available viruses was carried out using MAFFT 7.526 (https://mafft.cbrc.jp/alignment/software/, accessed on October 30, 2024) with multiple aa sequence alignments using G-INS-i (LbSV, LbCaV) and E-INS-i (LbPV, LbCV) as the best-fit model, respectively. The aligned aa sequences were used as input to generate phylogenetic trees through the maximum-likelihood method with the FastTree 2.1.11 tool available at http://www.microbesonline.org/fasttree/ (accessed on October 30, 2024). Local support values were calculated with the Shimodaira–Hasegawa test (SH) and 1,000 tree resamples. The polymerase proteins of Sin Nombre virus - NP_941976 and Thottapalayam virus - YP_001911124 (LbPV), Rice ragged stunt virus - AAC36456 (LbCV), Imperata yellow mottle virus - NC_011536, Southern cowpea mosaic virus - NC_001625 and Turnip rosette virus - NC_004553 (LbSV) were used as the outgroup in the phylogenetic trees.

## Results

### Virus discovery by data mining of Metatranscriptomic RNA-seq Libraries

To characterize the *L. botrana* virome, we analyzed existing RNA-Seq datasets deposited in the NCBI SRA from *L. botrana* individuals with diverse origins, biological characteristics and host conditions. Reads were retrieved from 29 publicly available libraries and processed. Through similarity searches against a viral protein database using BLASTX searches, we identified contigs and ORFs homologous to sequences from known viruses. Four novel viruses were discovered and reconstructed, and further characterized.

### Bioinformatics characterization of novel viral genomes A novel phasmavirus linked to L. botrana

A novel phasmavirus was identified and named Lobesia botrana phasmavirus (LbPV). This virus was detected in two transcriptome datasets from pheromone glands sampled from adult individuals collected in Germany in 2016 (**Table 1**). The reconstructed LbCPV genome (GenBank accession numbers BK067724-BK067726) comprised three segments (RNA1 = L, RNA2 = M and RNA3 = S) of single-stranded, negative-sense RNA of 6,458nt, 2,518nt and 1,759nt, respectively (**Fig. 1A**) with the 3’ end terminal sequence conserved among the viral segments, sharing the consensus sequence of other phasmavirus such as Anopheles triannulatus orthophasmavirus (AtoPV, **Fig. 1E**). Segment L has one ORF that encodes a putative RdRp protein of 2,100 aa; Segment M has one ORF that encodes a putative glycoprotein precursor of 766 aa (G); and the segment S has two overlapping ORFs, where ORF1 encodes a putative non-structural protein (NSs) of 125 aa, while the ORF2 encodes a putative nucleoprotein (NP) of 384 aa (**Fig. 1A**). Segment S was the most abundant one in terms of coverage, whereas the segment M was the one that accumulated less on those datasets where LbPV was identified (**Fig. 1A**). The RdRp protein shows the highest BlastP similarity with the moth-associated pink bollworm virus 2 (PBV2) RdRp, with 50.43% identity (**Supp. Table 1**), and included a Bunya_RdRp super family conserved domain (8C4V_A, bunyavirus RdRP, Hantaan virus, E-value 8e-104) identified in its sequence at aa positions 909-1,237 (**Fig. 1A**). All typical N-terminal domains, pre-motif and motifs A-E present in the bunyavirids encoded RdRps were identified in LBPV (**Fig. 1B**). The G protein showed the highest BlastP similarity with the moth-associated Seattle prectang virus (SEPV) G protein, with 32.63% identity (**Supp. Table 1**), and a EnvGly conserved domain (4HJ1_A ENVELOPE GLYCOPROTEIN; Class II fusion protein E-value 1.3e-21) was identified in its sequence at aa positions 392-466 (**Fig. 1A**). The putative G precursor is predicted to be processed by a conserved signal peptide peptidase to yield two mature G proteins (Gn and Gc) (**Fig. 1A** and **D**). The NP protein shows the highest BlastP similarity with the SEPV NP, with 43.56% identity (**Supp. Table 1**), and no conserved domains were identified in its sequence. The putative NSs has no hits in the BlastP searches, and no conserved domain was identified in its sequence, but it is worth mentioning that syntenic ORFs of analogous size and positions have been detected in other phasmavirids [15].

**Figure 1.**
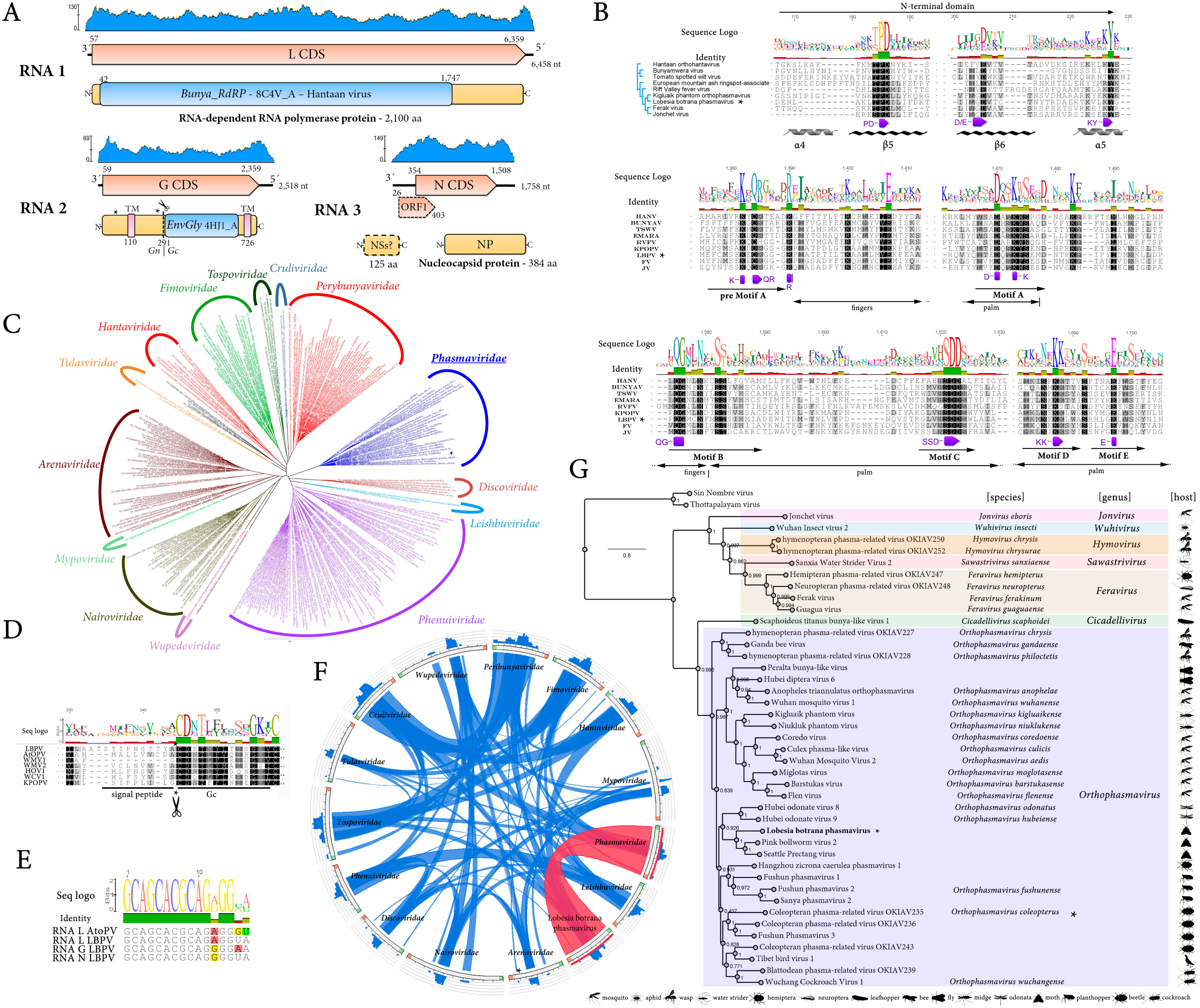
Genome organization, sequence conservation, phylogenetic relationships, and evolutionary insights into Lobesia botrana phasmavirus (LbPV) and related viruses. (**A**) Schematic representation of the genome organization of LbPV, showing its three segmented RNA genome. RNA 1 encodes the RNA-dependent RNA polymerase (RdRP), RNA 2 encodes the glycoprotein (G), and RNA 3 encodes the nucleocapsid protein (N). The annotations highlight functional domains, conserved motifs, and transmembrane regions (TM). Coding sequences are depicted in orange arrowed rectangles, predicted proteins in yellow rectangles and conserved domains in blue rectangles. The coverage plot (blue) displays the read depth across the genome and the number represents the maximum coverage. (**B**) Sequence conservation and alignment of the RdRP of LbPV and other bunyavirids. Sequence logos indicate conserved residues across related viruses. Functional motifs (e.g., Motif A, B, C, D, E) are labeled, and structural elements (e.g., α-helices and β-sheets) are indicated. (**C**) Circular phylogenetic tree showing relationships among members of *Phasmaviridae* and other viral families within the *Bunyavirales* order. Clades corresponding to different families are color-coded, and the tree highlights the evolutionary placement of LbPV within *Phasmaviridae*. (**D**) Sequence alignment of the glycoprotein cleavage site, showing conservation of the signal peptide and glycosylation motifs among LbPV and *Phasmaviridae* members. (**E**) Sequence logo of conserved terminal sequences of the 3′ RNA segments indicating base-pairing complementarity. (**F**) Circos plot depicting shared spatial protein domain architectures and sequence similarities across viral families within *Bunyavirales*. Connections are highlighted in blue (low similarity) to red (high similarity). (**G**) Maximum-likelihood phylogenetic tree of representative *Phasmaviridae* species, including members of genus *Orphophasmavirus* and closely related genera. The tree includes FastTree support values and host associations, with icons representing host types. Virus members of ICTV recognized species are indicated with binomial nomenclature. The scale bar represents the number of amino acid substitutions per site.

A phylogenetic tree based on LbPV RdRp and viruses belonging to different families within the class *Bunyaviricetes*, showed that LbPV is clustered with members belonging to the *Phasmaviridae* family, order *Elliovirales* (**Fig. 1C**). A Circus plot of spatial identity clearly supported a best complete coverage of LbPV only with phasmaviruses (**Fig. 1F**). A phylogenetic tree based on LbPV RdRp and viruses belonging to members of different genera within the *Phasmaviridae* family, showed that this virus is grouped with those viruses belonging to the *Orthophasmavirus* genus in a clade with the recently described moth-associated PBV2 and SEPV (**Fig. 1G**), showing a close evolutionary relationship of the moth-associated viruses within the *Orthophasmavirus* genus.

### A novel carmotetravirus linked to L. botrana

A novel carmotetravirus was identified and named Lobesia botrana carmotetravirus (LbCaV) (GenBank accession number BK067727). This virus was detected in ten transcriptome datasets corresponding to larval samples hosted by two different *V. vinifera* cultivars exposed to two different CO2 levels during two developmental stages (**Table 1**). The positive-sense, single stranded RNA genome of LbCaV is composed of 6,575 nucleotides (nt) and has three main ORFs (**Fig. 2A**). One ORF encodes the replicase fusion protein (p123) of 1,087 aa. This ORF contains a readthrough stop codon which results in the translation of a putative p58 protein of 521 aa or the fusion protein p123 (**Fig. 2A**). The p123 shows the highest BlastP similarity with the RdRp protein encoded by Hangzhou sesamia inferens carmotetravirus 1 (HSICTV1) with a 45.97% identity (**Supp. Table 1**) and includes a RDRP_SSRNA_POS conserved domain (Carmo_RDRP4, 8FMA_O RdRP, E-value 2.5e-18) in its sequence at aa positions 737-908 (**Fig. 2A**). The p58 protein shows the highest BlastP similarity with a hypothetical protein encoded by HSICTV1 with an 24.71% identity, and no conserved domains were identified in its sequence but two transmembrane regions (**Fig. 2A**). Another ORF, upstream of and overlapping with the replicase, encodes a putative protein p130 of 1,201 aa (**Fig. 2A**). This protein shows the highest BlastP similarity with a hypothetical protein encoded by HSICTV1 with an 27.49% identity (**Supp. Table 1**), and no conserved domain was identified in its sequence. The third main ORF, which is located downstream of that one encoding the replicase, encodes a putative coat protein (p87) of 796 aa (**Fig. 2A**). The CP shows the highest BlastP similarity with the CP protein encoded by the Providence virus (PrV), the member of the only International Committee on Taxonomy of Viruses (ICTV) recognized species within the single genus *Alphacarmotetravirus* of the family *Carmotetraviridae*, with a 50.51% identity (**Supp. Table 1**), and the Viral_coat conserved domain was identified in its sequence at aa positions 271-443 and 508-695 (**Fig. 2A**). Phylogenetic analysis based on the LbCaV replicase clustered this virus with a clade including the moth associated PrV and other putative members of the *Carmotetraviridae* family described on metagenomic studies, and away from the clades represented with members of the related *Permutetraviridae* and *Alphatetraviridae* families (**Fig. 2B**).

**Figure 2.**
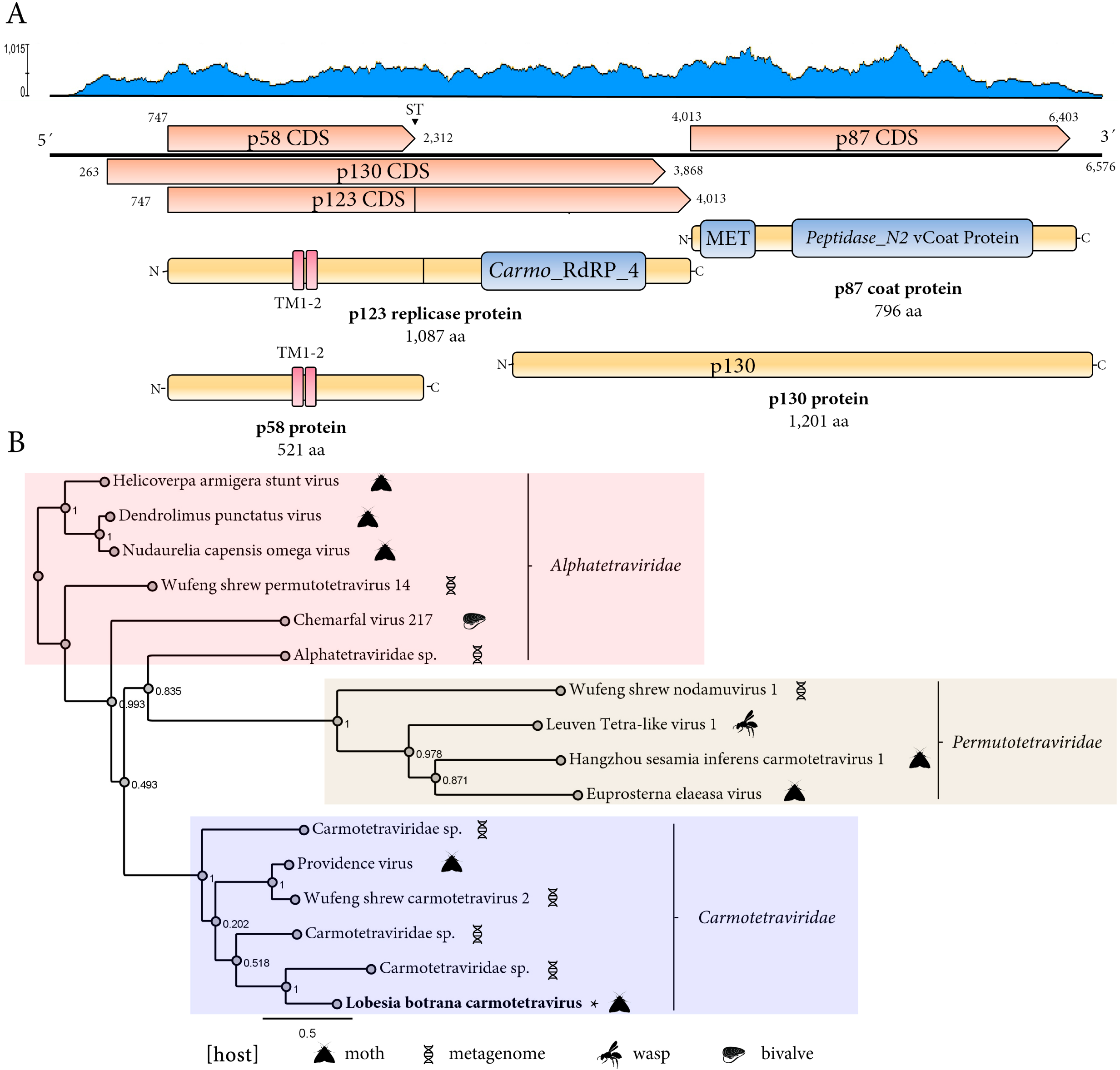
Genome organization and phylogenetic relationships of Lobesia botrana carmotetravirus (LbCaV) and related viruses. (**A**) Schematic representation of the genome organization of LbCaV. The linear genome is annotated with key coding sequences (CDS), including the replicase proteins (p123) the p130 and the coat protein (p87). Functional domains are highlighted, including Carmo_RdRP_4 (RNA-dependent RNA polymerase), MET (methyltransferase), and Peptidase_N2. Transmembrane regions (TM1-2) are indicated in the p58 protein, while protein sizes in amino acids (aa) are shown for major proteins. The coverage plot (blue) displays the read depth across the genome, with peaks corresponding to coding regions. Coding sequences are depicted in orange arrowed rectangles, predicted proteins in yellow rectangles and conserved domains in blue rectangles. (B) Maximum-likelihood phylogenetic tree of viruses from *Carmotetraviridae*, *Alphatetraviridae*, and *Permutotetraviridae*. Clades corresponding to each family are shaded (*Carmotetraviridae* in blue, *Alphatetraviridae* in pink, *Permutotetraviridae* in beige). Host associations are indicated with icons. FastTree support values are shown at nodes, and representative species are labeled. The scale bar represents the number of amino acid substitutions per site. This tree highlights the evolutionary relationships and host diversity among these virus families.

### A novel cypovirus linked to L. botrana

A novel cypovirus was identified and named Lobesia botrana cypovirus (LbCPV). This cypovirus was detected in 24 transcriptome datasets corresponding to larval samples hosted by two different *V. vinifera* cultivars exposed to two different CO2 levels during two developmental stages (**Table 1**). LbCPV genome is composed of ten double-stranded RNA (dsRNA) segments (**Fig. 3A**) (GenBank accession numbers BK067728-BK067737). The lengths of the ten segments range from 4,070 nt to 853 nt. Segment 1 length is 4,070 nt and has one ORF that encodes a putative protein of 1,331 aa. This protein shows the highest BlastP similarity with the Clanis bilineata cypovirus-type 23 (CbCPV-23) VP1 with an 88.43% identity (**Supp. Table 1**), and a Capsid protein VP1 3JB0_B conserved domain was identified in its sequence at aa positions 151-1,331. Therefore, this protein is the putative major capsid protein (**Fig. 3A**). Segment 2 length is 3,710 nt and has one ORF that encodes a putative protein of 1,183 aa. This protein shows the highest BlastP similarity with the Daphnis nerii cypovirus-type 23 (DnCPV-23) VP2 with an 87.15% identity (**Supp. Table 1**), and the CPV_RdRp_N, CPV_RdRp_pol_dom and CPV-RdRp_C Cypovirus 6K32_A conserved domains were identified in its sequence at aa positions 48-296, 327-697 and 847-1,179, respectively. Therefore, this protein is the putative RdRp protein (**Fig. 3A**). Segment 3 length is 3,308 nt and has one ORF that encodes a putative protein of 1,071 aa. This protein shows the highest BlastP similarity with the DnCPV-23 VP3 with an 84.59% identity (**Supp. Table 1**), and the 3JB0_A motif (9-1,069, E-value 2.7e-120) was detected including a Reov_VP3_GTase, Reov_VP3_MTase1 and Reov_VP3_MTase2 conserved domains in its sequence at aa positions 11-277, 483-703 and 872-1067, respectively. Therefore, this protein is a putative methyltransferase and guanylyltransferase structural protein (**Fig. 3A**). Segment 4 length is 3,781 nt and has one ORF that encodes a putative protein of 1,245 aa. This protein shows the highest BlastP similarity with the CbCPV-23 VP4 with an 79.87% identity (**Supp. Table 1**), and a PPPDE domain-containing protein; Cell attachment, Membrane penetration, VIRAL PROTEIN of Bombyx mori cypovirus 1 (7WHM_A, E-value 2e-102) was identified at aa positions 19-1,229. Thus, this protein is a putative minor capsid protein. Segment 5 length is 2,049 nt and has one ORF that encodes a putative protein of 642 aa. This protein shows the highest BlastP similarity with the DnCPV-23 VP5 with a 56.44% identity (**Supp. Table 1**), and no conserved domain was identified in its sequence (**Fig. 3A**). Its DnCPV-23 counterpart was suggested to be a putative structural protein [16]. Segment 6 length is 2,000 nt and has one ORF that encodes a putative protein of 612 aa. This protein shows the highest BlastP similarity with the DnCPV-23 VP6 with a 78.43% identity (**Supp. Table 1**), and no conserved domains. Its DnCPV-23 counterpart was suggested to be a protein with an unknown function [16]. Segment 7 length is 1,833 nt and has one ORF that encodes a putative protein of 537 aa. This protein shows the highest BlastP similarity with the CbCPV-23 VP7 with an 81.94% identity (**Supp. Table 1**), and a Viral structural protein 4 (6K32_B, E-value 1.1e-42); Cypovirus, VIRAL PROTEIN-RNA complex; HET: A2M conserved motif was identified at aa positions 13-530. Thus, this protein is a putative structural protein. Segment 8 length is 1,247 nt and has one ORF that encodes a putative protein of 391 aa. This protein shows the highest BlastP similarity with the DnCPV-23 VP8 with an 83.33% identity (**Supp. Table 1**), and a VP5 conserved domain (31Z3_D, E-value 8.7e-59) was identified in its sequence at aa positions 3-282. Thus, this protein is a putative structural protein. Segment 9 length is 1,105 nt and has one ORF that encodes a putative protein of 300 aa. This protein shows the highest BlastP similarity with the DnCPV-23 VP9 with a 79.26% identity (**Supp. Table 1**), and a NP protein; nucleocapsid protein, nucleoprotein, VIRAL PROTEIN motif was identified at aa positions 107-189 (4J4Y_D). Segment 10 length is 853 nt and has one ORF that encodes a putative protein of 246 aa. This protein shows the highest BlastP similarity with the DnCPV-23 VP10 with a 97.56% identity (**Supp. Table 1**), but the nt identity between LbCPV, DnCPV and CbCPV segment 10 is ca. 80%, which is in the range of the species demarcation criteria based on this segment for cypoviruses by the ICTV. Moreover, as expected, a CPV_Polyhedrin conserved domain (5A99_A, E-value 5.5e-53) was identified in its sequence at aa positions 3-244. Therefore, this protein is a putative polyhedrin protein (**Fig. 3A**). Phylogenetic analysis based on the LbCPV replicase placed this virus within the Lepidoptera, mostly moths, associated *Cypovirus* genus, clustering most closely with the recently described DnCPV-23 and CbCPV-23, suggesting the possibility that these 3 viruses could be members of a new species within genus *Cypovirus* (**Fig. 3B**).

**Figure 3.**
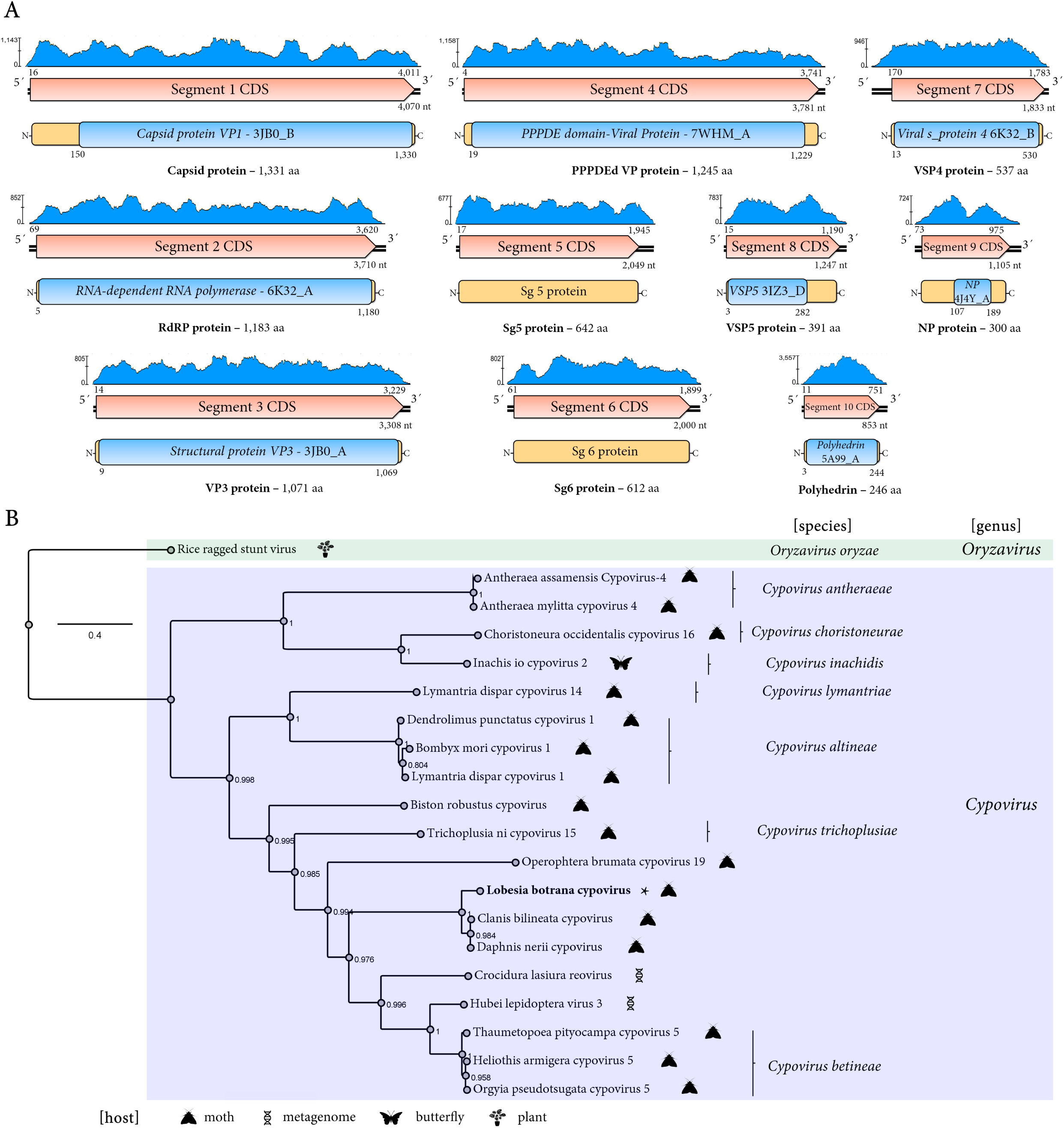
Genomic organization and phylogenetic analysis of Lobesia botrana cypovirus (LbCPV). (**A**) Genomic organization of LbCPV. The genome is segmented into 10 double-stranded RNA (dsRNA) segments (Seg 1-10). Each segment contains a single open reading frame (ORF) encoding a single protein. The predicted functions of the encoded proteins are indicated. The coverage plot (blue) displays the read depth across the genome. Coding sequences are depicted in orange arrowed rectangles, predicted proteins in yellow rectangles and conserved domains in blue rectangles. (**B**) Phylogenetic tree of LbCPV and cypoviruses based on the amino acid sequences of the replicase protein. The tree used as outgroup the orizavirus Rice ragged stunt virus. FastTree support values are shown at nodes, and representative species are labeled. The scale bar represents the number of amino acid substitutions per site. The hosts of the viruses are indicated by icons.

### A novel bi-segmented sobemo-like virus

A novel sobemo-like virus was identified and named Lobesia botrana sobemo-like virus (LbSV). This virus was detected in 27 transcriptome datasets including larval samples hosted by two different V. vinifera cultivars exposed to two different CO2 levels during two developmental stages, and whole adults susceptible or resistant to insecticides sampled in both Germany and Turkey (**Table 1**). The sequence of two strains LbSV_Ger (GenBank accession numbers BK067738-BK067739) and LbSV_Tur (GenBank accession numbers BK067740-BK067741) were assembled in this study. LbSV_Ger and LbSV_Tur genomes comprised two segments of single-stranded, positive-sense RNA (**Fig. 4A**). The RNA1 is composed of 2,701 nt and 2,622, respectively; while the RNA 2 comprises 1,413 and 1,379 nt, respectively (Fig. 4). LbSV_Ger and LbSV_Tur segment 1 has two ORFs, named as HP CDS and RdRP CDS. ORF HP encodes a hypothetical protein (HP) and an RdRP is encoded by the other and is expressed as a fusion polyprotein through a −1 ribosomal frameshift mechanism (**Fig. 4A**). The -1RFM signal consists of the slippery sequence 5′-GGGAAGC-3′ at coordinates 1,221-1,227. LbSV_Ger and LbSV_Tur HP have 459 aa and are 98% identical. This protein shows the highest BlastP similarity with Latepeofons virus (LtPV) with a 49.14% identity and 48.15% identity, respectively. A Peptidase_S1_PA conserved domain was identified in their sequence at aa positions 97-285 (Pro-VPg, 6FF0_A, **Fig. 4A**). LbSV_Ger and LbSV_Tur RdRp has 865 aa and are 98% identical. This protein shows the highest BlastP similarity with Wugcerasp virus 3 (WV3) with a 53.96% identity and 54.20% identity, respectively. An RNA-dir_pol-C was identified in their sequence (6QWT_A) at aa positions 510-851. LbSV_Ger and LbSV_Tur segment 2 has one ORF that encodes the CP with a size of 416 aa (**Fig. 4A**). Their CP are 98.3% identical and show the highest BlastP similarity with Buhirugu virus 16 (BHRGV16) CP protein with a 37.97% identity and 37.88% identity, respectively. A Nodavirus_capsid conserved domain (4WIZ_CX, E-value 2.9e-21) was identified in their sequence at aa positions 76-361 (**Fig. 4A**). Phylogenetic analysis based on the LbSV_Ger placed this virus within the clade containing unclassified solemovirids identified from a variety of arthropods, clustering most closely with the wasp associated WV3 (**Fig. 4B**).

**Figure 4.**
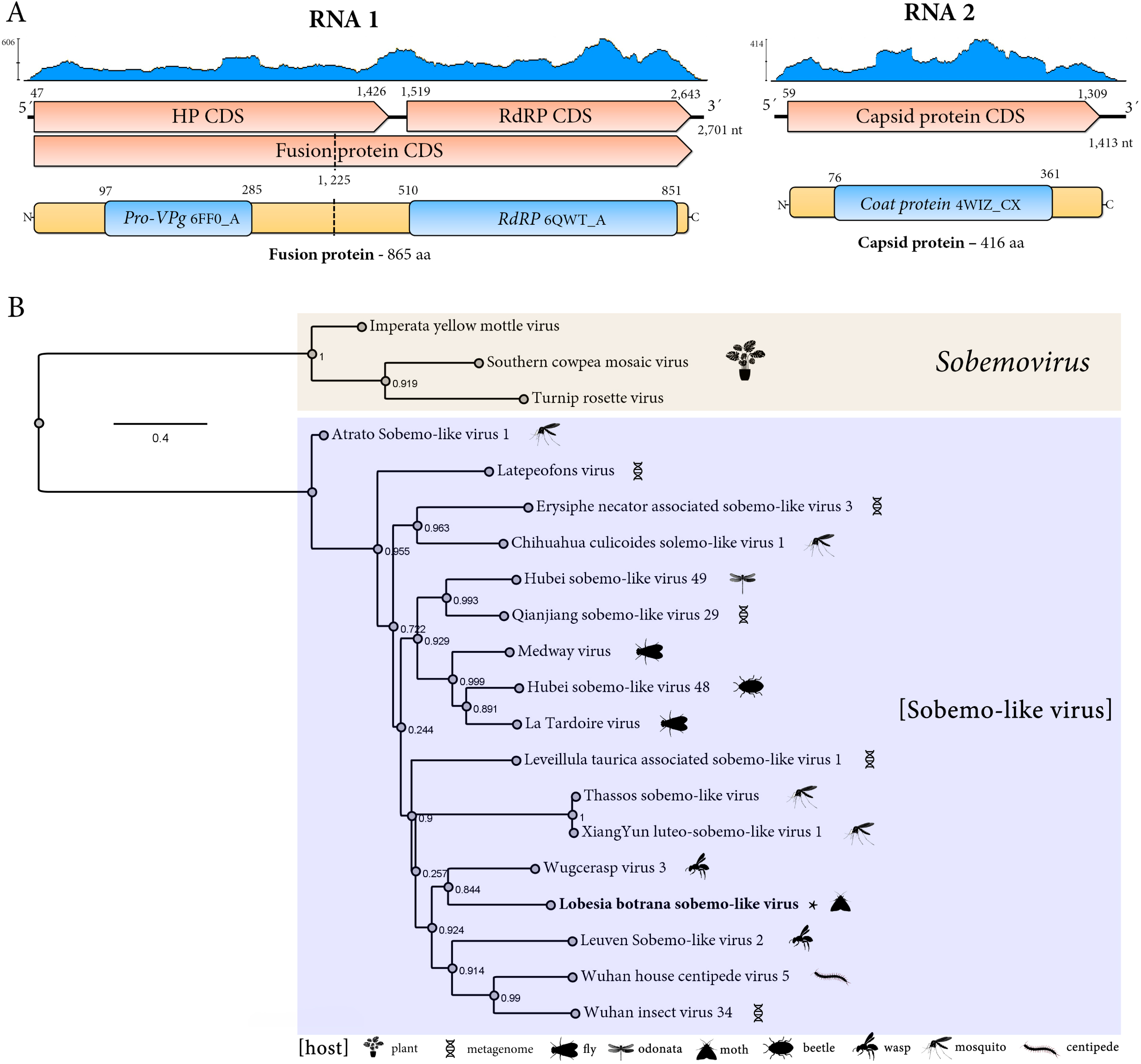
Genomic organization and phylogenetic analysis of Lobesia botrana sobemo-like virus (LbSV). (**A**) Genomic organization of LbSV. The genome is segmented into two double-stranded RNA (dsRNA) segments (RNA 1 and RNA 2). The predicted functions of the encoded proteins are indicated. The coverage plot (blue) displays the read depth across the genome. Coding sequences are depicted in orange arrowed rectangles, predicted proteins in yellow rectangles and conserved domains in blue rectangles. (**B**) Phylogenetic tree of LbSV, sobemovirus and sobemo-like viruses based on the amino acid sequences of the replicase protein. FastTree support values are shown at nodes, and representative species are labeled. The scale bar represents the number of amino acid substitutions per site. The hosts of the viruses are indicated by icons.

### A glimpse into expression level profiles and preliminary prevalence of L. botrana viruses

We investigated the presence and absence of specific RNA viruses in *Lobesia botrana* moths from Germany and Turkey. We analyzed both adults and larvae parasitizing Cabernet Sauvignon or Riesling grapevines at different growth stages, under conditions of high and low CO2 levels. Initial analyses revealed varying expression level profiles and prevalence of these viruses across different samples. The RNA expression levels of the four viruses associated with L. botrana—LbPV, LbSV, LbCPV, and LbCaV— were evaluated across the 29 RNA libraries, as shown in **Figure 5A**. The relative abundance of viral RNA, measured as fragments per kilobase of transcript per million mapped reads (FPKM), demonstrated marked variability across libraries. LbCPV was the most consistently and abundantly expressed virus across the libraries, followed by LbSV, LbCaV, and LbPV. Notably, certain libraries exhibited minimal or undetectable levels of viral RNA, reflecting a potential library-specific variability in viral presence or absence. **Fig. 5B** illustrates the RNA relative expression levels of the 10 genome segments of LbCPV. Each segment displayed distinct expression profiles, with segment 10—encoding the polyhedrin protein—showing the highest expression levels. This pattern aligns with the functional importance of polyhedrin in viral particle assembly [17]. While the expression of other segments was relatively uniform, minor variability among libraries suggests differential segment replication or transcript stability.

**Figure 5.**
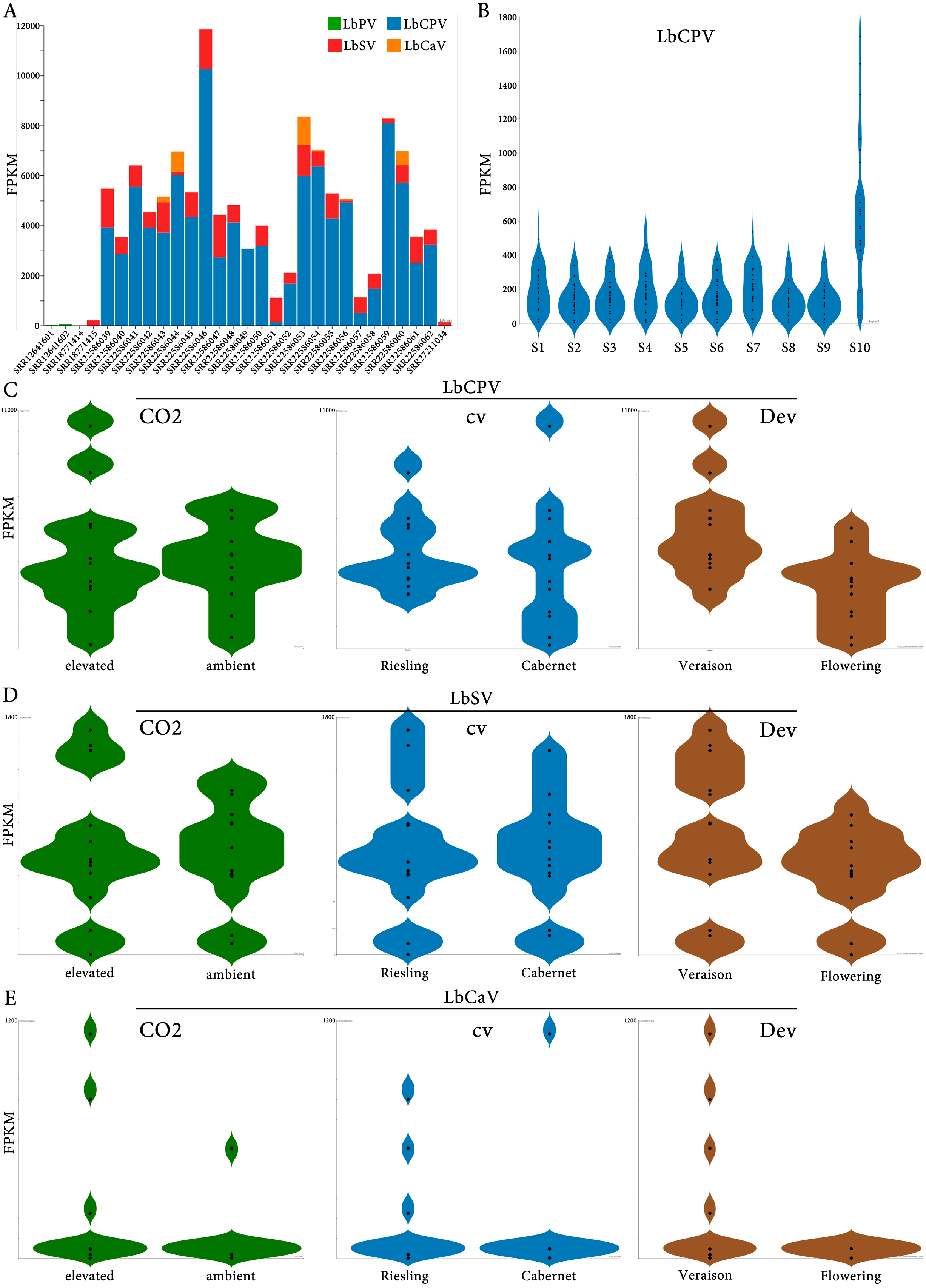
Assessment of relative RNA levels landscape of four Lobesia botrana-associated viruses in 29 RNA-seq secuencing libraries from various experimental conditions. (**A**) FPKM (Fragments Per Kilobase of transcript per Million mapped reads) values for four viruses associated with the grapevine moth, L. botrana: LbPV (L. botrana phasmavirus, green), LbSV (L. botrana sobemo-like virus, red), LbCPV (L. botrana cypovirus, blue), and LbCaV (L. botrana carmotetravirus, orange). (**B**) FPKM values for each of the 10 genome segments of LbCPV. Segment 10 encodes the polyhedrin protein, a structural component of the virus capsid. (C-E) FPKM values for LbCPV (**C**), LbSV (**D**), and LbCaV (**E**) in *Vitis vinifera* plants subjected to different environmental and developmental conditions: CO2: Plants were exposed to elevated (480 ppm) or ambient (400 ppm) CO2 levels. cv: moths were bred on two different grapevine cultivars: Cabernet Sauvignon and Riesling. Dev: moths were bred at two distinct developmental stages: flowering and veraison (onset of berry ripening). Each violin plot represents the distribution of FPKM values across multiple biological replicates. The width of the violin plot is proportional to the density of data points at that FPKM value.

**Fig. 5C, D**, and **E** present the RNA levels of LbCPV, LbSV, and LbCaV under varying environmental conditions, grapevine cultivars, and developmental stages of *V. vinifera*. Under elevated CO2 conditions, LbCPV exhibited slightly higher FPKM values compared to ambient CO2, suggesting that elevated CO2 may enhance viral RNA expression (**Fig. 5C**). In contrast, LbSV (**Fig. 5D**) and LbCaV (**Fig. 5E**) showed no significant differences in expression levels between the two CO2 conditions, suggesting the possibility that these viruses may be less responsive to atmospheric CO2 variations. Viral RNA levels were also compared between two grapevine cultivars: Cabernet Sauvignon and Riesling. For LbCPV, FPKM values were slightly higher in Riesling than in Cabernet Sauvignon, suggesting a potential host effect for this virus. Similarly, LbSV displayed marginally higher expression in Riesling, while LbCaV exhibited negligible differences between the two cultivars. The RNA expression of LbCPV varied between flowering and veraison stages, with higher levels observed during veraison (**Fig. 5C**). A similar trend was noted for LbSV (**Fig. 5D**). For LbCaV (**Fig. 5E**), RNA levels were lower overall, with only a minor increase during veraison, suggesting mild or no effect of cv host on RNA virus titters. Nevertheless, caution should be exercised when interpreting these RNA virus expression levels due to the relatively low sample size, which may limit the generalizability of the findings. Additionally, other unassessed variables, such as environmental stressors, microbial interactions, or host physiological conditions, may influence viral RNA levels and confound the observed patterns. Further investigations are warranted to elucidate the ecological and epidemiological factors driving the observed patterns and to assess the potential implications on the host.

## Discussion

### A glimpse into L. botrana viromics

Our high-throughput sequencing (HTS) based exploration of the *L. botrana* virome reveals a plethora of novel viral diversity. The discovered cypovirus lineage within *L. botrana* aligns with growing evidence highlighting host specialization and adaptation within Lepidoptera-associated viruses [18]. Notably, our analysis suggests potential inter-species transmission dynamics within *L. botrana* populations, echoing recent studies indicating horizontal virus transfer as a driving force in shaping insect viral communities and evolution [19, 20]. Further investigation into this cypovirus lineage, employing phylogenetic and comparative genomics approaches, will provide valuable insights into its evolutionary history and ecological roles within *L. botrana* populations.

### A novel phasmavirus

*Phasmaviridae* is a recently described family of negative-strand RNA viruses that are maintained in and/or transmitted by insects [21]. LbPV genome resembles that one reported for phasmavirids which possess a segmented genome, of three segments (L, M and S), that encode RNA-dependent RNA polymerase, a glycoprotein precursor and the nucleocapsid protein [21]. LbPV encoded RdRp have all the motifs identified in the bunyavirales encoded RdRps; which are essential for genome replication and transcription [22]. As was reported for the G precursor protein encoded by phasmavirids [21], the LbPV G precursor is processed by a conserved signal peptide peptidase to yield the mature Gn and Gc proteins which mediate cell entry of phasmavirions [23]. LbPV S segment encodes a putative NSs protein upstream of the NP, which was also found in some phasmavirids [21, 23], including the moth-associated PBV2 [24]. Moreover, the 3’ terminal nucleotides are conserved among the three LbPV segments and are likely complementary to the terminal sequences found in the 5’end which support the formation of panhandle-like structures that play a key role in RNA synthesis and genome packaging [25]. Conserved terminal segments among all three segments is characteristic of phasmavirids and it was suggested that the first seven of the terminal nucleotides may be genus specific [26].

LbPV represents the third reported phasmavirid identified in Lepidoptera, as this family was previously known to infect mostly mosquitoes and midges [21]. The three moth-associated phasmaviruses clustered together in the phylogenetic tree indicating a common evolutionary trajectory of these viruses suggesting a probable host-virus co-evolution. The BlastP searches as well as the phylogenetic analysis confirmed the placement of LbPV within the *Orthophasmavirus* genus. This virus should be classified as a novel member belonging to the *Orthophasmavirus* genus since the aa sequence identity of the L protein between the LBPV and related viruses is below 95%, which is the species demarcation criteria for this genus [21]. Unraveling the evolutionary trajectory and transmission dynamics of this novel phasmavirus within *L. botrana* demands further research, potentially elucidating novel ecological interactions within arthropod communities.

### A novel carmotetravirus

The novel carmotetravirus identified in this study, has a genome that exhibits the typical genomic organization of carmotetraviruses; which encode three main ORFs instead of the two ORFS that have other tetraviruses [27]. LbCaV, like the carmotetravirus PrV, has a replicase with a readthrough stop signal resulting in the production of an accessory protein. LbCaV replicase and accessory protein molecular weights are higher than that one reported for PrV [27]. The largest ORF, which overlaps the replicase, has a protein with similar size in LbCTV and PvR, and the presence, but with a low E-value, of a putative 2A-like processing sequence as the one located at the N terminus of PrV p130 [27]. Carmotetraviruses, belonging to the family *Carmotetraviridae*, possess a single-stranded, positive-sense RNA genome and only one genus, the *Alphacarmotetravirus*, has been created within this family, and its sole member PrV, has lepidopteran as its natural hosts [28]. Phylogenetic relationships grouped LbCTaV with PrV and some lepidopteran-associated proposed and recently described carmotetraviruses. Thus, LbCaV could be classified as novel member of a species within the *Alphacarmotetravirus* genus. The evolutionary history of PrV is not well understood, the expression of PrV is similar to that one observed in plant-infecting tombusviruses and PrV non-structural proteins are structurally related to plant tombusvirus and umbravirus accessory proteins [29]. [29] proposed that PrV is a hybrid virus with the potential to infect and replicate in both host plant and animal systems. This hypothesis was later confirmed by [30] who presented evidence that PrV can establish a productive infection in plants as well as in animal cells. Therefore, further studies should further characterize the biology and evolution of LbCaV.

### A novel cypovirus

The novel cypovirus identified in this study, has a genome that exhibits the typical segmented genomic organization of cypoviruses [31]. Cypoviruses, belonging to the family *Spinareoviridae*, represent a diverse and widely distributed group of segmented double-stranded RNA (dsRNA) viruses infecting a variety of arthropod hosts. So far, 16 different types of cypoviruses have been recognized by the ICTV, but many more cypoviruses are still to be recognized (https://ictv.global/report/chapter/spinareoviridae/spinareoviridae/cypovirus). Their segmented genome structure, a hallmark of the *Reovirales* order, offers unique features and flexibility in their evolutionary potential [31]. Each cypovirus genome typically consists of 10 dsRNA segments, each encoding one or two proteins crucial for viral replication, assembly, and pathogenesis. This modularity allows for intra- and inter-species reassortment, potentially contributing to the emergence of new viral strains and adaptation to diverse hosts [31]. Structural and functional annotation based on conserved domains identified features typical of cypoviruses in those LbCPV encoded proteins, including RdRp, capsid proteins, and virion assembly proteins which are essential components for viral replication, packaging, and assembly [32–36]. LbCPV is closely related to CbCPv-23 and DnCPV-23; however, it should be explored the possibility of being recognized as distinct species, rather than as a novel isolate of the cypovirus 23 type, since the nt identity for segment 10 is around the value set by the ICTV to demarcate species within the genus *Cypovirus* (https://ictv.global/report/chapter/spinareoviridae/spinareoviridae/cypovirus). DnCPV-23 was identified in the oleander hawk moth [16] while CbCPV-23 was identified in the soybean hawk moth [37]. Cypoviruses commonly induce chronic infection in the gut of insects [38] and have been suggested as promising agents for the control of lepidopteran species [39]. Interestingly, the closely related DnCPV-23 has a lethal effect on its host 16, 40]. Thus, LbCPV could be a promising virus for the control of *L. botrana*; therefore, further studies focused on the LbCPV effect on its host should be carried out.

### A novel sobemo-like virus

*Solemoviridae* is a recently established family of positive-strand RNA viruses which recognized members infect plants [41]. Typically, sobemovirids genomes are unsegmented with four to ten ORFs, which code for a viral suppressor of RNA silencing, a polyprotein (which contains a serine protease and a RdRp conserved domains) and a CP [41]. However, recently, sobemo-like viruses with two segments were identified in arthropods, such as crustaceans [42], myriapods [43] and insects [43–46]. Here we identified the sobemo-like virus LbSV, which adds to the group of sobemo-like viruses with bi-segmented genomes infecting non-plant hosts and is likely the first sobemo-like virus identified in moths. The genomic organization of segment 1 of LbSV and those bi-segmented sobemo-like viruses is highly similar to that one reported for the 5’ terminal half region of the sobemovirids (the ORF encoding the viral suppressor of RNA silencing protein is not present), because the first ORF encodes a polyprotein with a serine protease domain and the RdRp is translated via −1 programmed ribosomal frameshift (−1 PRF) from the next ORF [41]. LbSV slippery sequence 5′-GGGAAGC-3′ is highly similar to the one reported for poleroviruses, which is 5′-GGGAAAC-3′ [41]. Interestingly, LbSV CP, as was reported for other sobemo-like viruses CP [42] not only shares sequence similarity with other sobemo-like viruses, but also with noda-like viruses and permutotetra-like viruses. It has been hypothesized that because of capsid genes being encoded by subgenomic RNAs or genomic segments in these viruses there is a possibility that horizontal gene transfer could have been the outcome from mispackaging during co-infection of the same host [42]. Phylogenetic analysis based on the RdRp sequences placed LbSV within the clade containing unclassified sobemovirids. Thus, based on genomic organization and phylogenetic relationships at least a novel genus within the family *Solemoviridae* should be established to classify those solemo-like viruses which have arthropods as their hosts.

### An emergent landscape of the L. botrana virome

Beyond individual viral characterization, our analysis highlights the potential influence of the *L. botrana* virome on the moth-grapevine ecosystem. The genomic architecture of the novel viruses identified in this study provides valuable clues about their replicative strategies and evolutionary relationships. The segmented dsRNA genome of the cypovirus and the monosegmented carmotetravirus align in genomic organization with their established family counterparts, suggesting potential shared evolutionary pathways within these viral lineages. On the other hand, the genomic arrangement of the sobemo-like virus represent atypical features within their respective family, prompting further investigation into their replicative mechanisms and evolutionary origins.

Moving forward, our research opens exciting avenues for exploring the functional roles of these newly discovered viruses within *L. botrana*. Employing virus-host interaction studies and functional assays will unlock crucial insights into how these viruses influence their host’s physiology, behavior, and susceptibility to environmental factors. Understanding these interactions could pave the way for the development of novel pest control strategies, potentially targeting specific viral processes or manipulating virus-host interactions to interfere with the pest biology. The unveiled diversity of the *L. botrana*virome also calls for continued exploration of this hidden viral landscape. Advanced sequencing approaches, such as metagenomics and metatranscriptomics, hold immense potential for uncovering even greater viral diversity within *L. botrana* and other Lepidoptera species. Furthermore, functional screening techniques aimed at identifying virus-encoded virulence factors or host immunity pathways could offer valuable tools for elucidating the intricate interplay between *L. botrana* and its virome.

### Future Perspectives

While our initial investigation has unveiled the diversity of novel viruses residing within the European grapevine moth, a more profound understanding of their influence within this agricultural pest and the vineyard ecosystem necessitates further exploration. This necessitates deeper insights into the intricate relationship between virus and host, elucidating the specific roles these viruses play in shaping *L. botrana’s* biology and ecological interactions. Employing virus-host interaction studies using molecular and physiological techniques will allow us to observe the dynamic interplay between *L. botrana* and its holobiome, providing insights into how these viruses influence the moth’s metabolism, behavior, and resilience against environmental stressors.

The vineyard ecosystem is a complex web of interdependencies, and our newfound understanding of the *L. botrana* virome offers a glimpse into its hidden holobiome. This knowledge could pave the way for innovative biocontrol solutions that target vectors, disrupting the spread of plant diseases before they take root.

## Conclusions

Our high-throughput sequencing investigation has fostered our understanding of the European grapevine moth by unveiling a previously unseen viral world residing within this agricultural pest. The discovery of four novel viruses belonging to diverse families reflects a complex underexplored viral community potentially influencing *L. botrana* biology and impacting the broader vineyard ecosystem. The broad presence of these viruses in public metatranscriptomic datasets suggests their widespread prevalence and potential ecological significance. Additionally, the genomic architecture of each novel virus provides valuable clues about their replicative strategies and evolutionary relationships, opening doors for further exploration.

This research stands as a pivotal step in comprehending the *L. botrana* virome and its ecological implications. Unveiling this hidden dimension within *L. botrana* biology opens exciting avenues for sustainable pest management and a deeper understanding of the intricate viral webs shaping agricultural ecosystems. By embracing future research directions, we can harness the vast potential hidden within the *L. botrana* virome, ensuring a more sustainable future for vineyards and a broader understanding of the moth holobiome.

## Acknowledgments

We extend our sincere gratitude to the generators of the primary datasets utilized in this study. Their commitment to open science principles, evidenced by the availability of raw sequence data in public repositories, facilitated the generation of valuable insights through secondary data analyses. Their dedication to transparency and accessibility in scientific research has significantly contributed to the advancement of knowledge in our field.

## Author Contributions

Conceptualization, H.D and S.G.T.; data analysis, H.D and N.B; writing—original draft preparation, H.D; writing—review and editing, H.D, N.B, S.G.T. All authors have read and agreed to the published version of the manuscript.

## Institutional Review Board Statement

Not applicable for studies not involving humans or animals.

## Informed Consent Statement

Not applicable for studies not involving humans.

## Data availability statement

Nucleotide sequence data reported are available in the Third Party Annotation Section of the DDBJ/ENA/GenBank databases under the accession numbers TPA: BK067724 - BK067741. The virus sequences are also included as supplementary material for this submission.

