## Supplementary material for "RNA virus discovery sheds light on the virome of a major vineyard pest, the European grapevine moth (*Lobesia botrana*)": Supp. Table 1

**Supplementary Table 1**. Best BLASTP hits of each predicted gene products of *L. botrana* RNA virus sequences. Abbreviations: Query cover, blastp query coverage; na, not applicable; hp, hypothetical protein.

| **Genome segment/ protein** | **segment length (nt)** | **gene product (aa)** | **Description of best BLASTP hit** | **Score** | **Query cover** | **E-value** | **Identity** | **Accession** |
| --- | --- | --- | --- | --- | --- | --- | --- | --- |
| Lobesia botrana phasmavirus RNA1 | 6458 | 2100 | RNA-dependent RNA polymerase [Pink bollworm virus 2] | 2096 | 99% | 0.0 | 50.43% | QID77675.1 |
| Lobesia botrana phasmavirus RNA2 | 2518 | 766 | glycoprotein precursor [Seattle Prectang virus] | 395 | 92% | 7,00E-123 | 32.63% | YP_009666958.1 |
| Lobesia botrana phasmavirus RNA3 | 1759 | 384 | nucleocapsid [Seattle Prectang virus] | 273 | 84% | 2,00E-85 | 43.56% | YP_009666960.1 |
| Lobesia botrana carmotetravirus | 6576 | na | na | na | na | na | na | na |
| p58 translation | na | 521 | hp [Hangzhou sesamia inferens carmotetravirus 1] | 238 | 75% | 3,00E-64 | 24.51% | UHK03308.1 |
| p87 translation | na | 796 | capsid protein precursor [Providence virus] | 621 | 85% | 0.0 | 50.51% | AMQ67164.1 |
| p123 translation | na | 1087 | RNA-d RNA polymerase [Hangzhou sesamia inferens carmotetravirus 1] | 469 | 48% | 2,00E-150 | 45.97% | UHK03309.1 |
| p130 translation | na | 1201 | hprotein FSICV1_gp1 [Hangzhou sesamia inferens carmotetravirus 1] | 88.2 | 26% | 4,00E-13 | 27.49% | UHK03307.1 |
| Lobesia botrana cypovirus segment 1 | 4070 | 1331 | Cypovirus VP1 [Clanis bilineata cypovirus] | 2494 | 100% | 0.0 | 88.43% | UGZ05876.1 |
| Lobesia botrana cypovirus segment 2 | 3710 | 1184 | RNA-dependent RNA polymerase [Clanis bilineata cypovirus] | 2173 | 99% | 0.0 | 86.81% | UGZ05875.1 |
| Lobesia botrana cypovirus segment 3 | 3308 | 1071 | VP4 protein [Daphnis nerii cypovirus] | 1944 | 100% | 0.0 | 84.59% | YP_009551582.1 |
| Lobesia botrana cypovirus segment 4 | 3781 | 1245 | VP2 protein [Daphnis nerii cypovirus] | 1318 | 63% | 0.0 | 79.87% | UGZ05873.1 |
| Lobesia botrana cypovirus segment 5 | 2049 | 642 | putative structural protein P61 [Daphnis nerii cypovirus] | 761 | 100% | 0.0 | 56.44% | YP_009551580.1 |
| Lobesia botrana cypovirus segment 6 | 2000 | 612 | hypothetical protein VP6 [Daphnis nerii cypovirus] | 1058 | 100% | 0.0 | 78.43% | YP_009551578.1 |
| Lobesia botrana cypovirus segment 7 | 1833 | 537 | structural protein VP7 [Clanis bilineata cypovirus] | 954 | 100% | 0.0 | 81.94% | UGZ05870.1 |
| Lobesia botrana cypovirus segment 8 | 1247 | 391 | hypothetical protein [Daphnis nerii cypovirus] | 690 | 99% | 0.0 | 83.33% | YP_009551577.1 |
| Lobesia botrana cypovirus segment 9 | 1105 | 300 | hypothetical protein [Daphnis nerii cypovirus] | 508 | 99% | 7,00E-180 | 79.26% | YP_009551576.1 |
| Lobesia botrana cypovirus segment 10 | 853 | 246 | VP10 protein [Daphnis nerii cypovirus] | 508 | 100% | 0.0 | 97.56% | ARS43577.1 |
| Lobesia botrana sobemo-like virus segment 1 | 2701 | na | na | na | na | na | na | na |
| fusion protein translation | na | 865 | RNA-dependent RNA polymerase [Wugcerasp virus 3] | 520 | 52% | 2,00E-173 | 53.96% | WVD52809.1 |
| hypothetical protein 1 translation | na | 459 | hypothetical protein [Latepeofons virus] | 326 | 75% | 2,00E-104 | 49.14% | WNT71144.1 |
| RNA-dependent RNA polymerase | na | 374 | RNA-dependent RNA polymerase [Wugcerasp virus 3] | 466 | 99% | 1,00E-159 | 57.95% | WVD52809.1 |
| Lobesia botrana sobemo-like virus segment 2 | 1413 | na | na | na | na | na | na | na |
| capsid protein translation | na | 416 | capsid protein [buhirugu virus 16] | 291 | 97% | 3,00E-90 | 37.97% | WPV03041.1 |
