## Supplementary material for "RNA virus discovery sheds light on the virome of a major vineyard pest, the European grapevine moth (*Lobesia botrana*)": Table 1

**Table 1**. *Lobesia botrana* high-throughput RNA sequencing library datasets assessed for virus discovery. The runs are publicly available at the NCBI-SRA archive. Abbreviations: virusRPM, total virus reads per million reads in the specific run; LbCPV, Lobesia botrana cypovirus RPM; LbSV, Lobesia botrana sobemo-like virus RPM; LbPV, Lobesia botrana phasmavirus RPM; LbCaV, Lobesia botrana carmotetravirus RPM.

| **Run** | **BioProject** | **BioSample** | **Bases** | **SRA Study** | **virusRPM** | **LbCPV** | **LbSV** | **LbPV** | **LbCaV** |
| --- | --- | --- | --- | --- | --- | --- | --- | --- | --- |
| SRR12641601 | PRJNA663283 | SAMN16125021 | 11.76 G | SRP282412 | **26** | **0** | **0** | **26** | **0** |
| SRR12641602 | PRJNA663283 | SAMN16125021 | 11.22 G | SRP282412 | **62** | **0** | **0** | **62** | **0** |
| SRR18771414 | PRJNA827155 | SAMN27603370 | 7.76 G | SRP370512 | **8** | **0** | **8** | **0** | **0** |
| SRR18771415 | PRJNA827155 | SAMN27603369 | 10.09 G | SRP370512 | **211** | **0** | **211** | **0** | **0** |
| SRR22586039 | PRJNA910346 | SAMN32123568 | 3.52 G | SRP412145 | **5486** | **3925** | **1544** | **0** | **17** |
| SRR22586040 | PRJNA910346 | SAMN32123567 | 3.19 G | SRP412145 | **3533** | **2859** | **674** | **0** | **0** |
| SRR22586041 | PRJNA910346 | SAMN32123566 | 3.58 G | SRP412145 | **6403** | **5547** | **856** | **0** | **0** |
| SRR22586042 | PRJNA910346 | SAMN32123565 | 3.73 G | SRP412145 | **4537** | **3920** | **617** | **0** | **0** |
| SRR22586043 | PRJNA910346 | SAMN32123564 | 3.23 G | SRP412145 | **5152** | **3714** | **1213** | **0** | **226** |
| SRR22586044 | PRJNA910346 | SAMN32123563 | 3.71 G | SRP412145 | **6954** | **6008** | **145** | **0** | **801** |
| SRR22586045 | PRJNA910346 | SAMN32123562 | 4.16 G | SRP412145 | **5347** | **4331** | **997** | **0** | **18** |
| SRR22586046 | PRJNA910346 | SAMN32123582 | 3.78 G | SRP412145 | **11846** | **10261** | **1583** | **0** | **1** |
| SRR22586047 | PRJNA910346 | SAMN32123581 | 4.17 G | SRP412145 | **4428** | **2728** | **1699** | **0** | **1** |
| SRR22586048 | PRJNA910346 | SAMN32123580 | 3.85 G | SRP412145 | **4825** | **4125** | **700** | **0** | **0** |
| SRR22586049 | PRJNA910346 | SAMN32123579 | 3.27 G | SRP412145 | **3072** | **3071** | **1** | **0** | **0** |
| SRR22586050 | PRJNA910346 | SAMN32123561 | 3.57 G | SRP412145 | **3995** | **3187** | **808** | **0** | **0** |
| SRR22586051 | PRJNA910346 | SAMN32123578 | 3.08 G | SRP412145 | **1117** | **138** | **979** | **0** | **0** |
| SRR22586052 | PRJNA910346 | SAMN32123577 | 3.89 G | SRP412145 | **2108** | **1677** | **431** | **0** | **0** |
| SRR22586053 | PRJNA910346 | SAMN32123576 | 3.68 G | SRP412145 | **8353** | **5980** | **1243** | **0** | **1131** |
| SRR22586054 | PRJNA910346 | SAMN32123575 | 4.14 G | SRP412145 | **7016** | **6360** | **610** | **0** | **46** |
| SRR22586055 | PRJNA910346 | SAMN32123574 | 4.49 G | SRP412145 | **5283** | **4293** | **990** | **0** | **0** |
| SRR22586056 | PRJNA910346 | SAMN32123573 | 3.48 G | SRP412145 | **5058** | **4926** | **84** | **0** | **48** |
| SRR22586057 | PRJNA910346 | SAMN32123572 | 5.20 G | SRP412145 | **1131** | **498** | **633** | **0** | **0** |
| SRR22586058 | PRJNA910346 | SAMN32123571 | 3.34 G | SRP412145 | **2077** | **1474** | **603** | **0** | **0** |
| SRR22586059 | PRJNA910346 | SAMN32123570 | 3.48 G | SRP412145 | **8370** | **8093** | **277** | **0** | **0** |
| SRR22586060 | PRJNA910346 | SAMN32123569 | 3.96 G | SRP412145 | **6978** | **5703** | **721** | **0** | **554** |
| SRR22586061 | PRJNA910346 | SAMN32123560 | 4.31 G | SRP412145 | **3555** | **2497** | **1058** | **0** | **0** |
| SRR22586062 | PRJNA910346 | SAMN32123559 | 3.67 G | SRP412145 | **3832** | **3239** | **593** | **0** | **0** |
| SRR27211034 | PRJNA1050165 | SAMN38726487 | 436.05 M | SRP478043 | **147** | **0** | **147** | **0** | **0** |
